# Lysosomal permeabilization by Group A *Streptococcus* releases proteins into the macrophage cytosol

**DOI:** 10.1101/2025.10.25.684498

**Authors:** Ava Quezada, Kevin Lord, Cheldon Alcantara, Claire Delahunty, Kevin Kim, Olivia Okamoto, John R Yates, Cheryl YM Okumura

**Affiliations:** Biology Department, Occidental College, Los Angeles, CA, 90041 USA; Department of Integrative Structural and Computational Biology, Scripps Research Institute, La Jolla, CA 92037, USA

**Keywords:** Group A Streptococcus, lysosome, macrophage, streptolysin O (SLO), IL- 1β, histones, inflammasome

## Abstract

The human-specific bacterial pathogen Group A *Streptococcus* (GAS) is a significant cause of morbidity and mortality due to its ability to cause severe invasive infection. Although macrophages are important for controlling GAS infection, we and others have demonstrated that GAS can persist in macrophages by perforating the phagolysosome using the pore-forming toxin streptolysin O (SLO). In this study, we examined how phagosomal perforation releases lysosomal and bacterial proteins into the cytosol and alters cytosolic protein content in the macrophage. Using IL-1β as a measure of intracellular pathogen detection, we confirmed that cytosolic preparations from macrophages infected with either wild-type (WT) or SLO-deficient (ΔSLO) bacteria contained new proteins that are absent in uninfected cytosol controls. Proteomic analysis revealed distinct cytosolic protein profiles in both WT- and ΔSLO-infected macrophages. M1 protein was detected only in the cytosol of WT-infected macrophages and corresponded with the IL-1β response, indicating SLO-mediated release of M1 protein from the phagosome, and providing a mechanism for cytosolic recognition of this virulence factor. Unexpectedly, cytosolic extracts of both WT- and ΔSLO-infected macrophages contained all histone proteins, suggesting that nucleosomal complexes are released into the cytosol during GAS infection. Our work reveals both a mechanism for the activation of the inflammatory response on a cellular level, and the surprising consequences of phagosomal perforation in GAS infections. These responses may collectively contribute to the pathologies observed during severe invasive GAS infection, and can help inform therapies aimed at improving macrophage function and patient outcomes.

**Importance:** The post-pandemic increase in cases of invasive Group A Streptococcus (GAS) infection, along with its sudden onset and severity of symptoms, underscores the need to understand GAS survival in host cells. Macrophages are key first responder immune cells essential for limiting the spread of GAS infection. As professional phagocytes, macrophages typically engulf bacteria in phagosomes and digest them using proteolytic enzymes. However, the human-specific pathogen GAS has acquired several evasion tactics through co-evolution with our immune system. In previous work, we have shown that GAS survives digestion by perforating the phagosome. In this study, we determined that this perforation allows both host and bacterial proteins to leak into the cytosol. These leaked proteins can dictate the subsequent response of the macrophage and may contribute to the pathology observed in invasive infection. Understanding this response can lead to improved therapies for GAS infections.

## Introduction

In addition to millions of annual cases of non-invasive infections, Group A *Streptococcus* (GAS; *Streptococcus pyogenes*) can cause severe invasive GAS (iGAS) infections such as cellulitis, bacteremia, pneumonia, streptococcal toxic shock syndrome and necrotizing fasciitis. In 2023, as part of a post-pandemic increase in iGAS infection, there were an estimated 43,100 life-threatening infections in the United States (1,2). A similar rise in iGAS infection was seen in Europe, concurrent with the rise and spread of M1UK GAS (3–6). Severe inflammation is associated with iGAS, particularly in necrotizing fasciitis (7), and increased inflammation markers in plasma have been observed in pediatric iGAS cases (8). Although GAS continues to be susceptible to beta-lactam antibiotics, efficient identification and treatment of iGAS, where disease can progress quickly, remains a challenge (9).

Macrophages are key first responder immune cells essential for limiting the spread of GAS infection from soft tissues (10–12). In a human challenge trial of pharyngitis, monocytes, the precursor cell of macrophages, were the dominant responsive cell type and mediated a strong pro-inflammatory response (13). In their role as professional phagocytes, macrophages quickly engulf pathogens in phagosomes, and lysosomal fusion delivers proteolytic enzymes to facilitate bacterial destruction. We and others have shown that GAS, a human-specific pathogen that co-evolved with our immune system, has multiple mechanisms for surviving this macrophage response.

These include 1) perforating the lysosomal membrane using the pore-forming toxin Streptolysin O (SLO) or recruiting the host protein CD63/LAMP-3, which releases ions and lysosomal proteins into the cytosol, and 2) inhibiting v-ATPase activity, preventing lysosomal acidification (14–16). Following lysosomal permeabilization, GAS remains in lysosomes, where it neither replicates nor escapes into the cytosol, enabling persistence in a cell that normally functions to kill them (7,12,16,17).

While GAS-mediated perforation of the phagolysosomal membrane allows bacterial survival, it also permits the release of lysosomal contents into the macrophage cytosol. In our previous work, we demonstrated that SLO causes the leakage of proteins up to 40kD in size into the cytosol, including active cathepsin B, a lysosomal enzyme (16). The presence of cathepsin B in the cytosol acts as a danger signal that can trigger a secondary host response such as activation of the NLRP3 inflammasome, resulting in the proteolytic processing and secretion of IL-1β (18,19). This secondary response in macrophages may serve as an important safety net mechanism to facilitate the clearance of non-functional macrophages and recruit additional immune cells for pathogen clearance. It is well-established that GAS induces NLRP3 inflammasome activation, and this activation is proposed to be protective in invasive infections (20,21). In previous reports, the GAS M1 surface protein has been shown to stimulate NLRP3 activation (22). Additionally, the cysteine protease SpeB can directly cleave pro-IL-1β in an NLRP3-independent manner to generate an inflammatory response (21,23). In both cases, bacterial proteins must be delivered intracellularly to interact with their targets, but the mechanism of this delivery to the cytosol remains unclear. Furthermore, it is unknown whether the concentrations of these GAS proteins in an individual cell are sufficient to trigger this response. SpeB is a secreted protein that does not accumulate to appreciable levels until GAS are in stationary phase (24).

In this study, we isolated cytosol from infected macrophages to identify both host and bacterial proteins released during GAS macrophage infection and phagolysosomal permeabilization. While we used IL-1β response as a measure of macrophage second-line responses to infection, our data reveal that GAS infection induces unexpected changes in the cytosolic protein profile, introducing or eliminating proteins that may influence cellular response. These results inform our understanding of bacterial persistence and host response, providing a foundation for designing effective treatment strategies for iGAS infections.

## Methods

### Antibodies and chemicals

LPS derived from *Salmonella enterica*, recombinant SLO and CA-074 methyl ester (CA074) were purchased from MilliporeSigma. Antibodies to the following proteins were used in this study: LAMP-1 (Cell Signaling Technology, #9091), LAMP-2 (Abcam, ab25631), Cathepsin B (Cell Signaling Technology, D1C7Y, #31718), Histone H3 (Cell Signaling Technology, D1H2, #4499), Lamin A/C (Cell Signaling Technology, 4C11, #4777) and GAPDH (Santa Cruz Biotechnology, 0411 sc-47724).

### Bacterial strains

Wild-type (WT) GAS strain M1T1 5448 (M1 GAS) was originally isolated from a patient with necrotizing fasciitis and toxic shock syndrome (25). Isogenic mutant strains lacking the *slo* (ΔSLO) and *emm1* (ΔM1) genes were previously described (26,27). Precise allelic replacement of the genes was sequence verified for each strain. Group B *Streptococcus* (GBS) WT strain serotype III (COH1) was originally isolated from a neonate with early onset sepsis (28). Bacterial strains were cultivated in Todd-Hewitt broth (THB) at 37°C. For all experiments, bacteria were grown to log phase in the presence of 1:200 pooled normal human serum (Thermo Fisher Scientific) to opsonize bacteria. Heat-killed (HK) bacteria were prepared by incubating a known concentration of bacteria at 95°C for 10 min, followed by a 15-min opsonization in 1:200 pooled normal human serum at room temperature.

### Cell culture

THP-1 cells were purchased from Sigma (cat. 88081201) and cultured in RPMI supplemented with 10% heat-inactivated fetal bovine serum (FBS) (Cytiva), 2mM L-glutamine and 100 U/mL penicillin/100 µg/mL streptomycin at 37°C/5% CO2. Cells were differentiated to macrophages using 20-40nM phorbol 12-myristate 13-acetate (PMA) (MilliporeSigma) 48 hours prior to experiments.

### Cytosol isolation and Western blot

Log phase bacteria were prepared in RPMI supplemented with 2% FBS (infection medium) and added to 10^7^ macrophages at a multiplicity of infection (MOI) of 10. Plates were centrifuged at 480 x g for 5 minutes to initiate and synchronize bacterial contact with cells. Cells were co-cultured with bacteria for 1 hour at 37°C, 5% CO2. Both culture supernatants (containing detached cells) and attached cells were collected and washed twice with ice-cold PBS. To selectively lyse the plasma membrane (29), cells were incubated with 30 µg/mL digitonin in acetate buffer (50 mM Na-acetate pH 5.6, 150 mM NaCl, 0.5 mM EDTA) supplemented with protease inhibitors (HALT protease inhibitor cocktail, ThermoFisher, 100 µM PMSF, 1 µg/mL pepstatin) and 0.1 µg/mL penicillin for 1 hour on ice. Cell lysates were vortexed at maximum speed for 30 seconds and centrifuged at maximum speed (18,400 x g) for 15 minutes at 4°C to pellet membrane proteins. The pellet consisting of intact organelles including lysosomes, was resuspended in acetate buffer and collected as the membrane fraction. The supernatant (cytosol) was filtered through a 0.22 µm filter syringe to exclude contaminating bacteria. Cytosolic fractions were centrifuged in 5 kDa molecular weight cut-off (MWCO) columns at 12,000 x g to concentrate samples (Fig. 1C) or 30 kDa MWCO columns at 15,000 x g for 10 minutes at 4°C to separate the content by size (all other data). All fractions were stored at -80°C. Protein concentrations were measured by BCA assay (Thermo Fisher Scientific) and 10ug of each sample was run on SDS-PAGE gels and transferred to a 0.2 um PVDF membrane. Blots were blocked with 5% (w/v) non-fat dry milk and probed with the indicated antibodies overnight at 4°C.

**Fig. 1:**
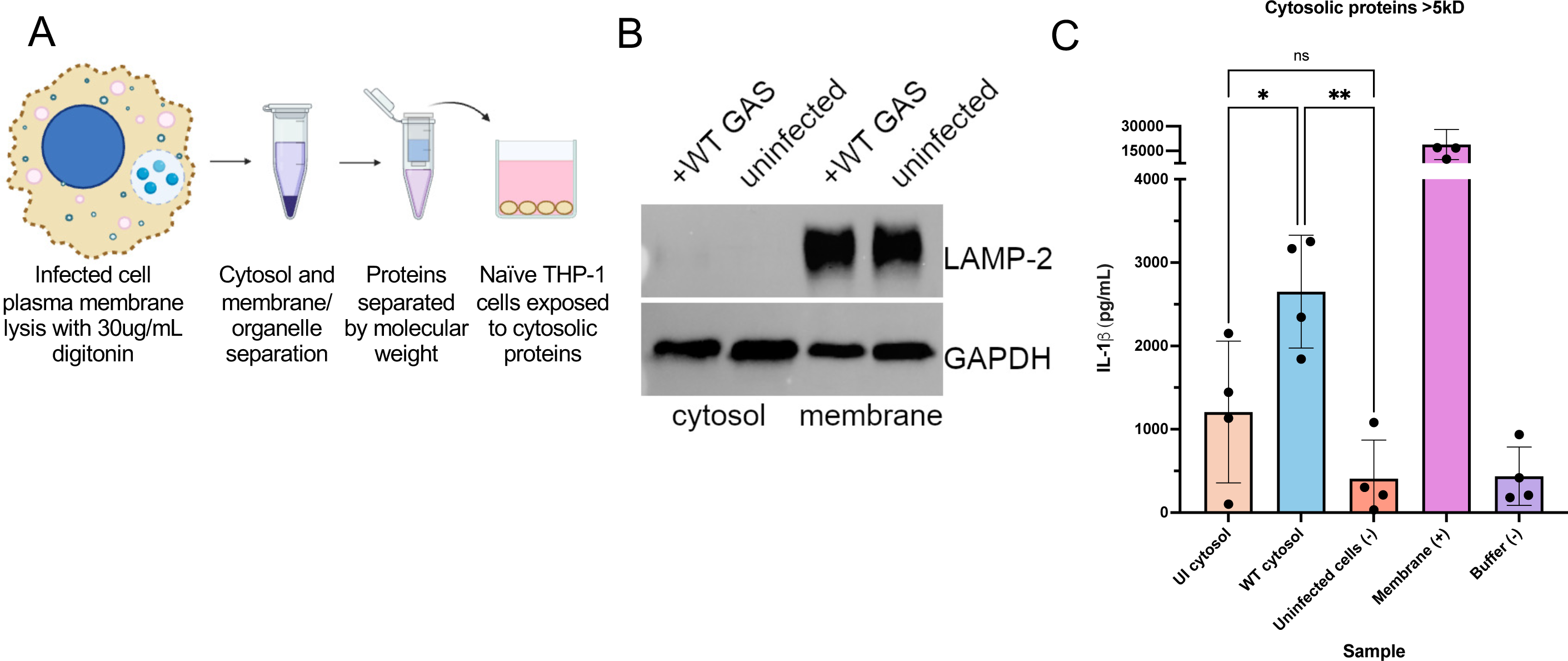
Proteins in GAS-infected cell cytosol stimulate IL-1β secretion. **A)** Cytosolic protein isolation schematic. Image was created with Biorender (biorender.com). **B)** Cytosolic and membrane fractions from WT GAS-infected and uninfected cells were probed by Western blot for the presence of LAMP-2 (lysosomal membrane protein) and GAPDH (cytosolic protein). **C)** Cytosolic fractions were tested for inflammasome- activating capacity by exposing naïve THP-1 cells to cytosolic fractions from uninfected (UI cytosol) or WT GAS-infected (WT cytosol) cells and measuring IL-1β secretion by ELISA. THP-1 cells incubated in media only (uninfected cells), with membrane fractions or fractionation buffer were included as controls. Experiments were performed with at least 3 different cytosolic protein preparations and ELISAs were performed in duplicate for each sample. Data are represented as mean ± SD and analyzed by one-way ANOVA with Tukey’s multiple comparison (*p<0.05, **p<0.01, ns = not significant).

### ELISA

5 x 10^5^ THP-1 cells were infected with the indicated live bacteria at an MOI of 10 in infection medium for 1 hr. For infection experiments with cytosolic fractions, cytosol fractions were adjusted in cell culture media (RPMI+10% FBS) to 20ug per 5 x 10^5^ THP- 1 cells and exposed to cells for 24 hrs. Culture supernatants were collected and centrifuged to pellet detached cells and cell debris. Release of human IL-1β into the culture supernatant was determined by enzyme-linked immunosorbent assay (ELISA) (R&D Systems) in accordance with the manufacturer’s instructions. Plates were read at 450 nm with a plate reader. IL-1β was calculated in each treatment in reference to the provided standard. All samples were measured in duplicate and experiments were repeated at least 3 independent times.

### Label-free proteomics analysis

Samples were MeOH/CHCl3 precipitated and the pellet was air-dried. Pellets were resuspended in 60 uL of buffer (8M urea, 100mM Tris, pH 8.5) and reduced with 3 uL of 100 mM tris(2-carboxyethyl) phosphine hydrochloride (TCEP). Samples were alkylated in the dark for 20 min with 250 mM iodoacetamide and digested with trypsin overnight at 37°C. Samples were acidified with formic acid (5%), and 1 ug of sample was loaded onto EvoTips (Evosep) according to the manufacturer’s protocol. Samples were run on an Evosep One (Evosep) coupled to a timsTOF Pro mass spectrometer (Bruker Daltonics). Peptides were separated with a gradient of buffer A (0.1% formic acid in H2O) and buffer B (0.1% formic acid in acetonitrile) on a 15 cm × 150 μm ID column with BEH 1.7 μm C18 beads (Waters) and an integrated tip pulled in-house. MS scans were acquired in PASEF mode, with one MS1 TIMS-MS survey scan and ten PASEF MS/MS scans per 1.1 s acquisition cycle. Both ion accumulation time and ramp time in the dual TIMS analyzer were set to 100 ms, and the ion mobility range was 1/K0 = 0.6 to 1.6 V s cm−2. The m/z range was 100−1700. Precursor ions selected for MS/MS analysis were isolated with a 2 Th window for m/z < 700 and a 3 Th window for m/z >700. Collisional energy was lowered linearly from 59 eV at 1/K0 = 1.6 V s cm−2 to 20 eV at 1/K0 = 0.6 V s cm−2 as a function of increasing mobility. Precursors for MS/MS were picked at an intensity threshold of 2500, a target value of 20,000, and an active exclusion of 24 s. Singly charged precursor ions were excluded with a polygon filter.

Tandem mass spectra were extracted from raw files using RawExtract (Version 1.9.9) and were searched against a concatenated Human/Streptococcus pyogenes database combined_UniProt_NCBI_Human_Streptococcus_pyogenes_03-01-2023 with reversed sequences using ProLuCID. A static modification of carbamidomethylation on cysteine (57.02146) was considered. Data were searched with 50 ppm precursor ion tolerance and 600 ppm fragment ion tolerance. Data were filtered using DTASelect2 to a protein false-positive rate of <1%. A minimum of two peptides per protein and one tryptic end per peptide were required. Statistical models for peptide mass modification (modstat) and tryptic status (trypstat) were applied.

### Proteomic data analysis

Independent preparations of uninfected (UI, 4 replicates), WT-infected (WT, 4 replicates) and ΔSLO-infected (SLO, 3 replicates) cytosolic fractions were analyzed. Contaminants including keratin and reverse hits were removed from the data sets before analysis. The data sets were filtered to only include proteins that were identified in at least two of the biological replicates for each sample to yield a list of 1532 proteins that were confidently identified (Supp. Fig. 1, Supp. Table 1). Normalized spectral abundance factors (NSAF) were log2 transformed and normalized by scaling each value against the average of all the proteins within the sample, followed by normalization by correlation slope (30). missForest was used to impute missing values in replicate samples (31) and values below the least expressed protein were imputed for proteins consistently absent in a sample type. Principal component analysis (PCA) was performed using prcomp in R on individual replicates using squared cosine values. Heat maps were generated using heatmaply (32). Proteins of interest were analyzed using ShinyGO 0.82 (33).

### DNA quantification

Total DNA was quantified via a Qubit 3.0 fluorometer using the dsDNA HS Assay Kit (ThermoFisher Scientific). 2ul of cytosolic fraction or digitonin acetate buffer (control) was diluted in Qubit dsDNA HS reagent/buffer solution. DNA was measured against the provided standard and calculated relative to the amount of protein in the sample. Digitonin acetate buffer spiked with THP-1 genomic DNA was also included as a control (data not shown). All samples were measured in duplicate and experiments were repeated at least 3 independent times.

### Statistics

All graph data were analyzed using Prism v. 10.6.0 (GraphPad Software), by one-way ANOVA with Tukey’s multiple comparison tests. Outliers were assessed by the ROUT method (Q = 1%). For all data presented, ****P<0.0001, ***P<0.001, **P<0.01, *P<0.05, n.s., not significant.

## Results

### Cytosolic fractions from WT GAS-infected cells have inflammasome-stimulating activity

In our previous work, we demonstrated that GAS infection resulted in lysosomal permeabilization and leakage of large lysosomal proteins into the cytosol, including cathepsin B which can cause activation of the NLRP3 inflammasome (16,19). To determine which lysosomal and bacterial proteins were released into the cytosol, as well as to determine global cytosolic protein changes in response to infection and lysosomal leakage, we developed a cytosolic protein isolation protocol for proteomic analysis (Fig. 1A). Cells were infected for 1 hour with the indicated bacteria to allow lysosomal permeabilization and release of lysosomal contents into the cytosol. Cells were lysed with 30ug/mL digitonin, which lyses the plasma membrane, but keeps organellar membranes intact (29). Cell lysates were spun to separate cytosolic proteins from membrane proteins and intact organelles, then treated with antibiotics and filtered with a 0.2um filter to eliminate the presence of live bacteria in the sample (data not shown).

Samples were concentrated in columns with a 5kD molecular weight cut-off and analyzed by Western blot (Fig. 1B). The absence of the lysosomal membrane protein LAMP-2 confirmed the resulting cytosolic fractions were free of intact lysosomes (Fig. 1B). GAPDH was found in high abundance in the cytosolic fraction as expected, and was also present in the membrane fraction since not all cytosol was collected to prevent disturbance of the membrane pellet (Fig. 1B). We next tested whether our cytosolic fractions had inflammasome-stimulating activity by exposing naïve PMA-differentiated THP-1 macrophages to cytosolic fractions and measuring IL-1β secretion (Fig. 1A). In previous work, stimulation and differentiation of THP-1 cells using PMA provides a sufficient priming signal to allow inflammasome activation by GAS M1 protein, while other cell types such as mouse bone marrow-derived macrophages require pre- stimulation with agonists such as LPS (22,34). Pre-stimulation with LPS caused increased basal IL-1β secretion in uninfected cells and a high IL-1β response to all cytosolic fractions regardless of infection status (Supplemental Fig. 1). We therefore tested untreated (no LPS pre-stimulation) PMA-differentiated THP-1 macrophage responses to cytosolic fractions (Fig. 1C). THP-1 macrophages released an increased amount of IL-1β in response to cytosolic fractions from WT-infected cells compared with cytosolic fractions from uninfected cells, confirming that we were capturing proteins in our cytosolic fraction that had inflammasome-stimulating activity (Fig. 1C). Thus, we were able to successfully isolate cytosolic fractions from GAS-infected cells that contained proteins that were leaked from permeabilized lysosomes and had inflammasome-stimulating activity.

### Cytosolic fractions from ΔSLO-infected cells also have inflammasome- stimulating activity

The pore-forming toxin SLO permeabilizes the phagolysosomal membrane to allow the release of >40kD proteins into the cytosol, including cathepsin B, which could stimulate inflammasome activity and IL-1β secretion (16,19). Others have demonstrated that SLO itself can stimulate inflammasome activity (20,34). We therefore tested whether the inflammasome-stimulating activity we observed from cytosolic fractions of WT GAS-infected cells (Fig. 1C) was due to the presence of SLO. We found that in the absence of bacteria, recombinant SLO alone did not have inflammasome-stimulating activity until added at high concentrations (Fig. 2A).

**Fig. 2:**
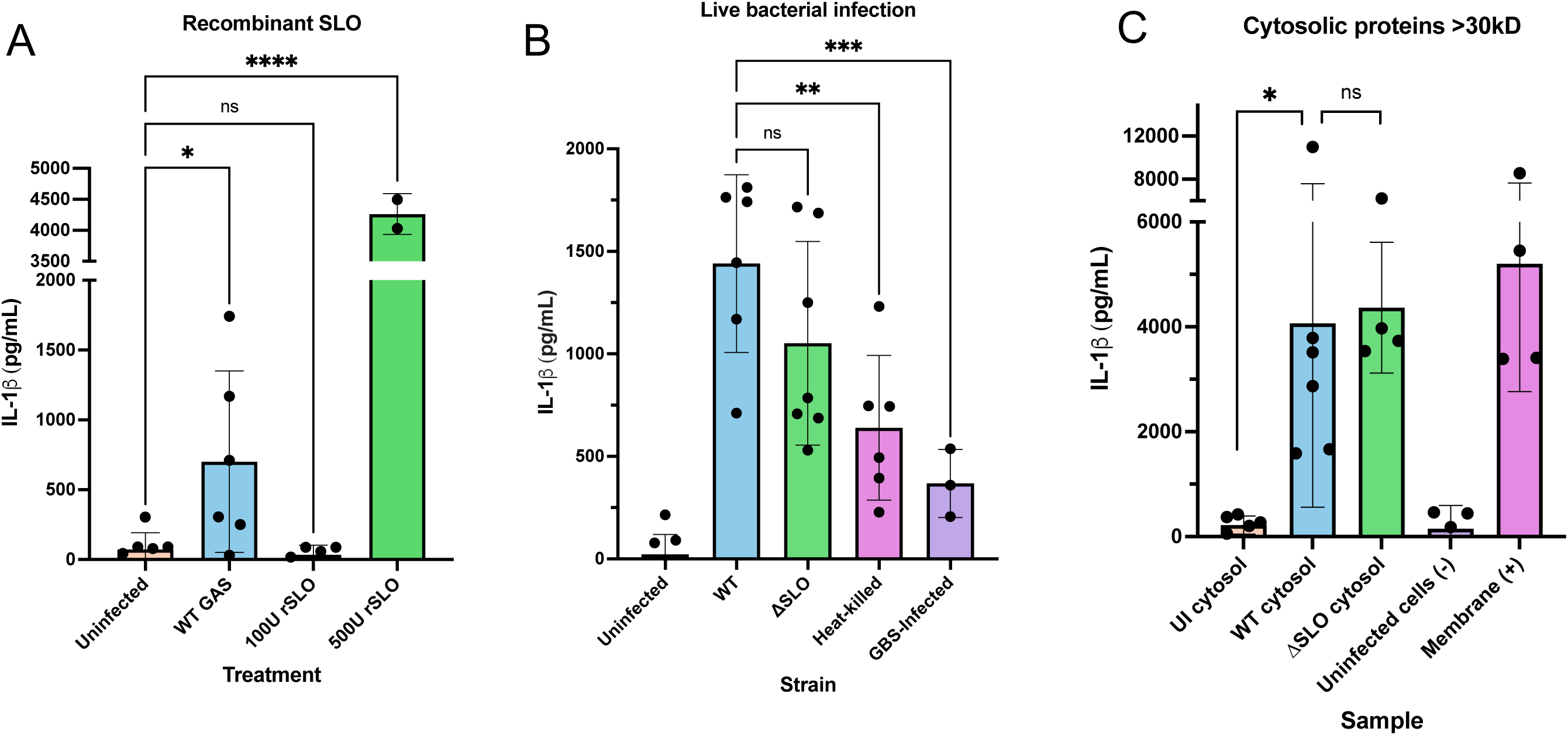
Cytosolic proteins from ΔSLO-infected cells also stimulate IL-1β secretion. IL-1β secretion was measured by ELISA for PMA-differentiated THP-1 macrophages responses to **(A)** recombinant SLO (rSLO), **(B)** live bacteria or **(C)** cytosolic proteins >30kD isolated from THP-1 cells infected with the indicated strains. Experiments were performed at least 3 independent times (A, B) or with at least 3 different cytosolic protein preparations (C) and ELISAs were performed in duplicate for each sample. Data are represented as mean ± SD and analyzed by one-way ANOVA with Tukey’s multiple comparison (*p<0.05, **p<0.01, ns = not significant).

We next infected THP-1 macrophages with the live ΔSLO mutant strain (Fig. 2B). Previous studies have demonstrated that mutant bacteria lacking SLO do not stimulate NLRP3 inflammasomes to the same extent as WT-infected cells (22,34). In contrast to previous findings, the inflammasome-stimulating activity was not significantly different between cells infected with WT GAS or the ΔSLO mutant in the aggregate (Fig. 2B), though in individual experiments infection with the ΔSLO mutant strain always produced a slightly lower IL-1β secretion response than the WT strain. These data indicated that there was still inflammasome-inducing activity caused by the ΔSLO mutant. We also infected cells with heat-killed WT GAS, which maintains surface proteins but is incapable of secreting bacterial proteins, including SLO. In agreement with previous results, infection with heat-killed bacteria, which does not cause lysosomal permeabilization, did not produce a notable IL-1β response (Fig. 2B)(16,20). Finally, we also infected cells with live Group B Streptococcus (GBS) and found a decreased IL-1β response compared with GAS infection, similar to other work (35). Thus, infection with live GAS caused an IL-1β response in THP-1 macrophages, regardless of SLO expression.

In our previous work, we found that cells infected with ΔSLO mutant bacteria also had permeabilized phagolysosomal membranes, mediated by CD63/LAMP-3 (16). We therefore tested cytosol from cells infected with the ΔSLO mutant strain for inflammasome-stimulating activity (Fig. 2C). For these experiments and for downstream proteomic analyses, to limit our analysis to larger proteins that are still capable of escaping the lysosome (16), we concentrated cytosolic fractions in 30kD molecular weight cut-off columns. WT-infected cytosol not only retained inflammasome-activating capacity (Fig. 2C) but also induced a stronger response than cytosolic fractions obtained using 5kD molecular weight cut-off columns (Fig. 1C). In agreement with our live bacterial infection (Fig. 2B), cytosol from ΔSLO-infected cells also elicited a strong IL-1β response (Fig. 2C). Collectively, our data indicate that GAS-induced permeabilization of phagolysosomes causes an IL-1β response, regardless of whether permeabilization is mediated by SLO or by other cellular mechanisms.

### Proteomic analysis of cytosolic fractions reveals proteins released by GAS permeabilization of the phagolysosome

To analyze which proteins may be escaping GAS-permeabilized phagolysosomes, we next performed label-free proteomics analyses on cytosolic fractions isolated from uninfected, WT-infected, or ΔSLO-infected THP-1 macrophages. Across all samples, we confidently identified a total of 1532 proteins (Supp. Table 1).

PCA (Supp. Fig. 2D) and heatmap analyses (Fig. 3A) demonstrated that independent preparations of each sample gave reproducible protein profiles. GAS proteins were only found in infected cells as expected (Supp. Table 1 and 2). Cross-referencing the total protein list with GO terms for nuclear matrix (GO:0016363) resulted in a very small list of hit proteins that were not exclusive to the nucleus (Supp. Table 2), confirming that our cytosolic fractions were well-isolated and free of organellar contaminants. Notably, the largest differences between samples were the presence/absence of proteins, though there were some differences in expression levels (Fig. 3A, B). We therefore focused our analysis on proteins that were present/absent in our samples (Fig. 3B).

**Fig. 3:**
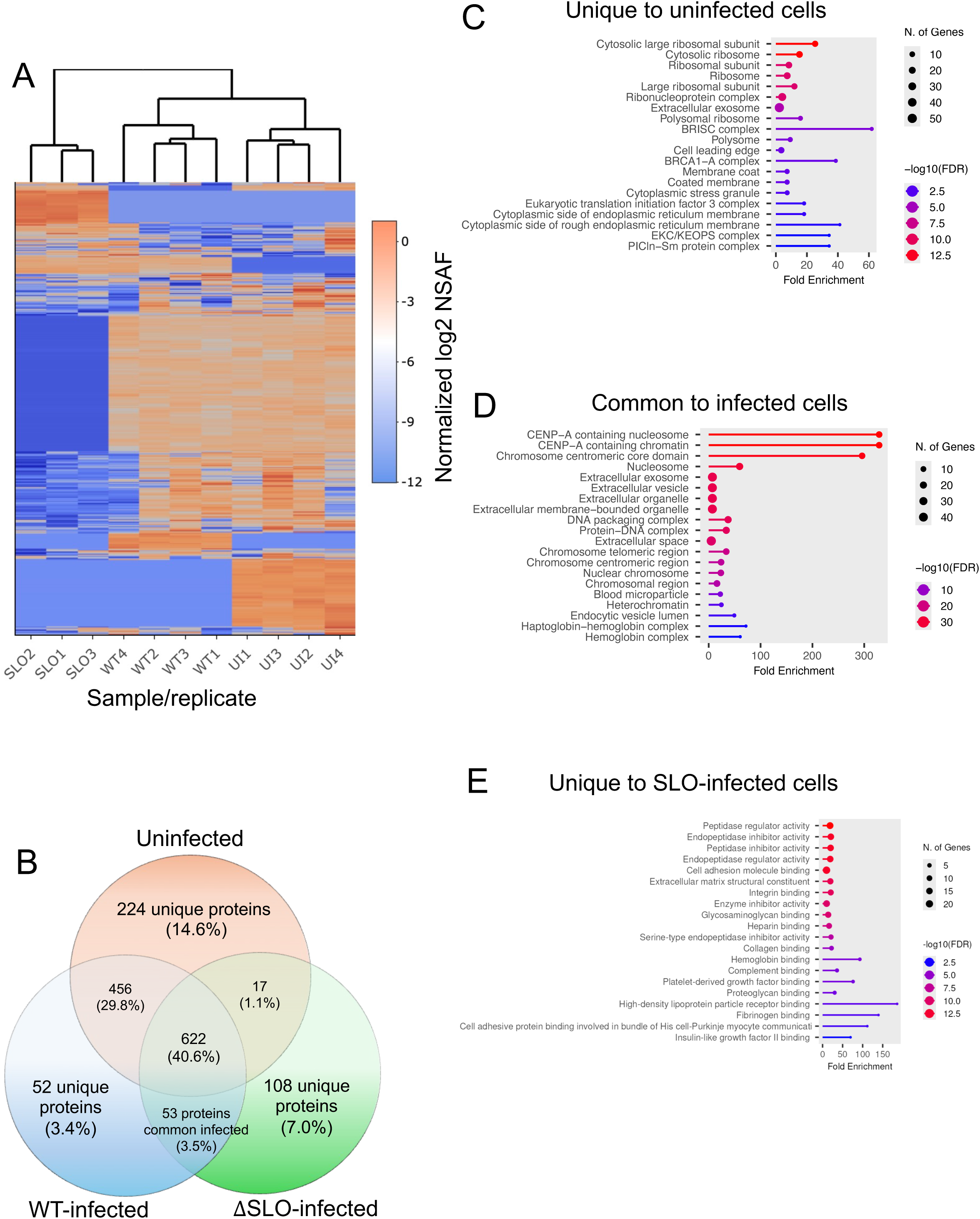
Proteomic analysis of cytosolic proteins from GAS-infected cells. (**A)** Relative abundance of proteins in replicate preparations of cytosol from uninfected (UI), WT- or ΔSLO-infected (SLO) cells. **(B)** Number of proteins identified in each sample. **(C-E)** Fold-enrichment analysis of proteins in indicated samples. Images were generated in ShinyGO 0.82.

Of the proteins found exclusively in uninfected cell cytosol, surprisingly about 15% were ribosomal subunits or ribosome-associated proteins (Fig. 3C, Supp. Table 2 and 3). Translation initiation factors, DnaJ proteins (molecular chaperones), and ubiquitin-conjugating enzymes were also present in uninfected, but not GAS-infected cytosolic fractions (Supp. Table 2 and 3), indicating that protein translation and folding may be negatively affected in GAS-infected cells. About 15% of proteins unique to uninfected cells were also associated with extracellular exosomes (Fig. 3C, Supp. Table 2 and 3).

In our previous work, we showed that infection with ΔSLO mutant bacteria also results in phagolysosomal permeabilization, supporting that GAS proteins can be found in the cytosol of ΔSLO-infected cells (Supp. Table 2). Common to cytosolic fractions from both WT- and ΔSLO-infected cells, we found secreted GAS proteins, including streptococcal GAPDH (Supp. Table 2). Proteins annotated as nicotine adenine dinucleotide glycohydrolase precursor and streptolysin O precursor were found in both WT- and ΔSLO-infected cells (Supp. Table 2). This annotation may reflect that the proteomic analysis detected the inhibitor of the NADase (encoded by *ifs*) and/or the NADase protein since the ΔSLO mutant was created with precise allelic replacement and retains expression of the other genes in the *ifs-slo-nga* operon (26). Most surprisingly, we found almost all the histone proteins in the cytosol of GAS-infected cells (Fig. 3D, Supp. Tables 2 and 3). In addition to the histones, many proteins were also classified as extracellular exosome or vesicle proteins (Supp. Tables 2 and 3). Although some of the histone proteins were cross-listed as extracellular exosome proteins, almost half of the proteins common to both WT- and ΔSLO-infected cells were associated with the extracellular exosome. Combined with the data for proteins found exclusively in uninfected cells, this may indicate that GAS infection significantly alters exosome/extracellular vesicles, either in formation or content.

For proteins exclusively found in the cytosol of WT-infected cells, we found lysosomal lumen proteins, as expected given that we have demonstrated SLO pores can induce leakage of dextrans >40kD (Supp. Table 2 and 3)(16). These proteins included cathepsin B, which we had previously reported to be present in our cytosolic fractions, and lysosomal glycosidases (16). Interestingly, we consistently found M1 protein in WT-infected, but not in ΔSLO-infected cells (Supp Table 2), which is likely due to the different-sized pores created by SLO in WT-infected cells or CD63/LAMP-3 in ΔSLO-infected cells.

For proteins exclusively found in ΔSLO-infected cells, we found extracellular matrix (ECM) or ECM-binding proteins, including several collagen proteins and integrins (Fig. 3E, Supp. Table 2 and 3). Immunoglobulin proteins were also present in high abundance (Supp. Table 2), though it is unclear whether this was in part due to opsonizing the bacteria with human serum prior to infection. Finally, we found a number of proteins associated with the complement pathway, including C3, C4, C5, C7, and C8, and complement factors B and H (Supp. Table 2 and 3). Additional GAS proteins were uniquely found in the ΔSLO-infected cytosolic fractions, including the cysteine protease SpeB (Supp. Table 2). Our results indicate that phagosomes in ΔSLO-infected cells can be permeabilized, but likely due to the differences in pore size, release different proteins and induce a distinct cellular response from WT-infected cells.

### WT GAS-infected cells allow release of M1 protein into the cytosol to stimulate an IL-1β response

In previous work, recombinant M1 protein added to PMA-stimulated THP-1 macrophages could stimulate IL-1β activation and secretion (22). However, it was unclear how M1 could be introduced into the cytosol of macrophages during natural infection. Given that we found M1 protein in the cytosol of WT-infected cells (Supp. Table 2), we hypothesized that M1 protein released from WT-permeabilized phagolysosomes could be responsible for the IL-1β secretion we observed (Fig. 1C, 2C). We therefore isolated cytosolic proteins from THP-1 cells infected with the ΔM1 isogenic mutant and tested whether they could stimulate an IL-1β response (Fig. 4A). In contrast to cytosolic preparations from ΔSLO-infected cells (Fig. 2C), THP-1 cells had a significantly decreased IL-1β response to cytosolic preparations from ΔM1-infected cells (Fig. 4A). These results indicate that M1 protein released from WT GAS-permeabilized phagosomes accounts for the majority of the inflammasome activation we observe in response to WT-infected cytosol.

**Fig. 4:**
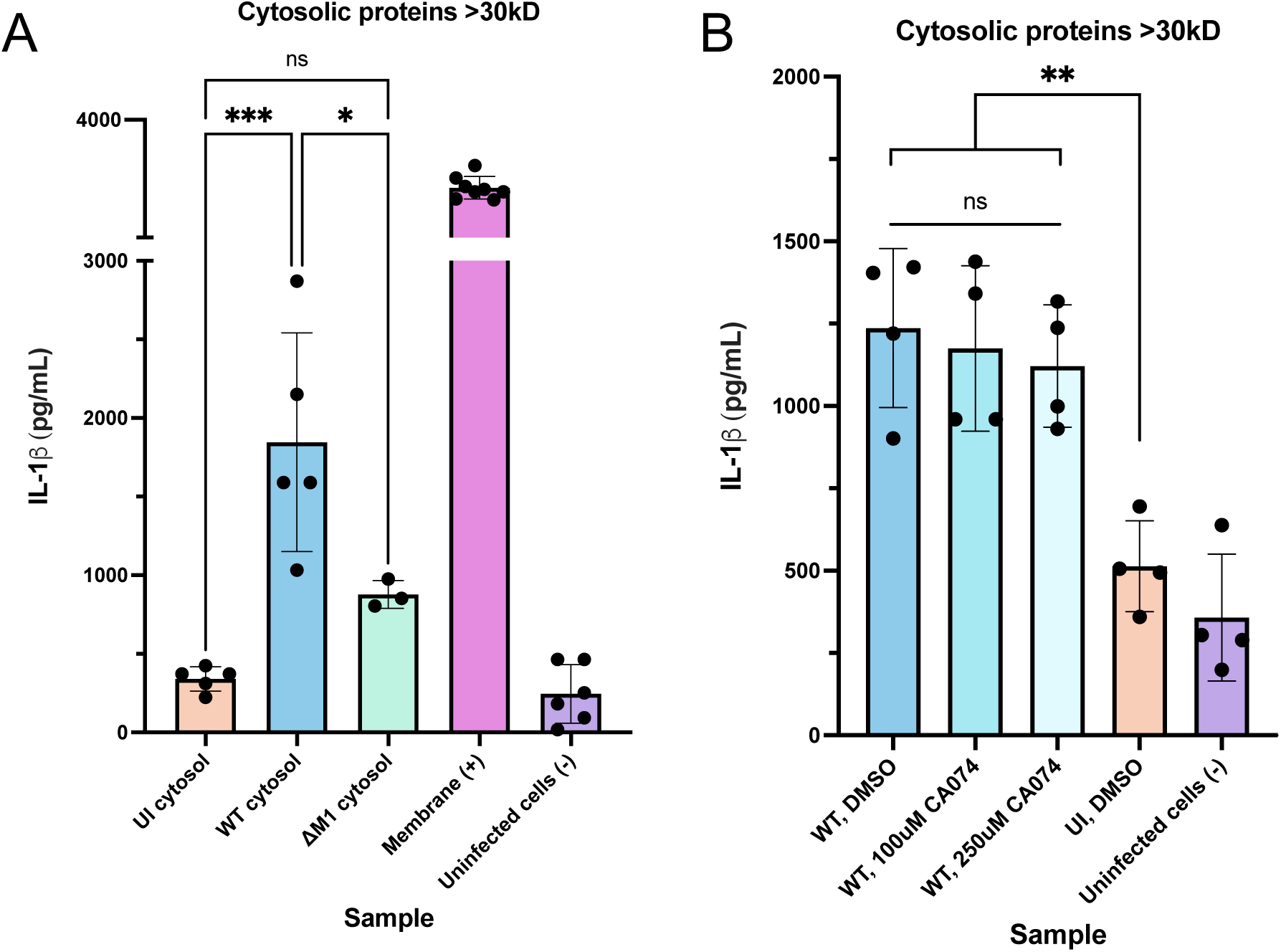
GAS M1 protein, but not lysosomal enzymes trigger an IL-1b response. **A)** Cytosolic fractions from uninfected (UI), WT-infected, or ΔM1-infected or cells were incubated with naïve THP-1 cells for 2 hours. **B)** Cells were pre-incubated with the indicated concentration of CA-074 Me or DMSO, then exposed to cytosolic proteins from WT-infected or uninfected (UI) cells for 2 hours. THP-1 cells incubated in media only (uninfected cells), with membrane fractions or fractionation buffer were included as controls as indicated, and IL-1β secretion was measured by ELISA. Experiments were performed with at least 3 different cytosolic protein preparations and ELISAs were performed in duplicate for each sample. Data are represented as mean ± SD and analyzed by one-way ANOVA with Tukey’s multiple comparison (*p<0.05, **p<0.01, ns = not significant).

Our previous work showed that cathepsin B released into the cytosol retains activity (16), and others have demonstrated that cathepsin B is capable of stimulating the NLRP3 inflammasome (19,36). Cathepsin B was not detectable by Western blot in our cytosolic preparations (Supp. Fig. 3), which may be due to cathepsin B being close to the molecular weight cut-off we used for concentrating the cytosolic fractions and reducing the amount of detectable cathepsin B in the sample. However, we consistently identified cathepsin B in the cytosol in both WT- and ΔSLO-infected cells (Supp. Table 2). We therefore tested the ability of our WT-infected cytosolic fractions to induce an IL- 1β response in the presence of CA-074 methyl ester (CA-074), a potent and specific cathepsin B inhibitor (36,37), and found no significant reduction in IL-1β response (Fig. 4B). Combined, our data suggest that M1 protein released from permeabilized phagolysosomes is primarily responsible for the IL-1β response observed in WT-infected cells.

### Histones are released into the macrophage cytosol in response to GAS infection

A surprising finding from our proteomic analysis was the identification of histones in the cytosol of both WT- and ΔSLO-infected cells (Fig. 3D, Supp. Table 2). The lack of contaminating organellar proteins, including nuclear matrix proteins, in our proteomic analysis suggests that release of these proteins is specific to GAS infection (Supp. Table 2). Given the small size of individual histone proteins (10-23kD), we hypothesize that full nucleosomes are released into GAS-infected cytosol since we used 30kD molecular weight cut-off columns in the cytosolic preparations and identified almost all histone subunits (Fig. 3D, Supp. Table 2 and 3). To confirm this finding, we performed a Western blot on our cytosolic fractions and looked for the presence of the histone H3 subunit, which was regularly present in WT- but not ΔSLO-infected cytosol (Supp. Table 1). Consistent with our findings in our proteomics analysis, we detected histone H3 in the cytosolic fractions of both WT- and ΔSLO-infected cells (Fig. 5A). We did not detect LAMP-1 or Lamin A/C, a nuclear membrane protein, confirming that our cytosolic preparations were not contaminated with organelles, especially nuclei (Fig. 5A). To determine whether histone presence in cytosolic fractions were specifically due to GAS infection, we also prepared cytosol from Group B Streptococcus (GBS)-infected cells. We confirmed that histones were only present in the cytosol of GAS-infected cells (Fig. 5B).

**Fig. 5:**
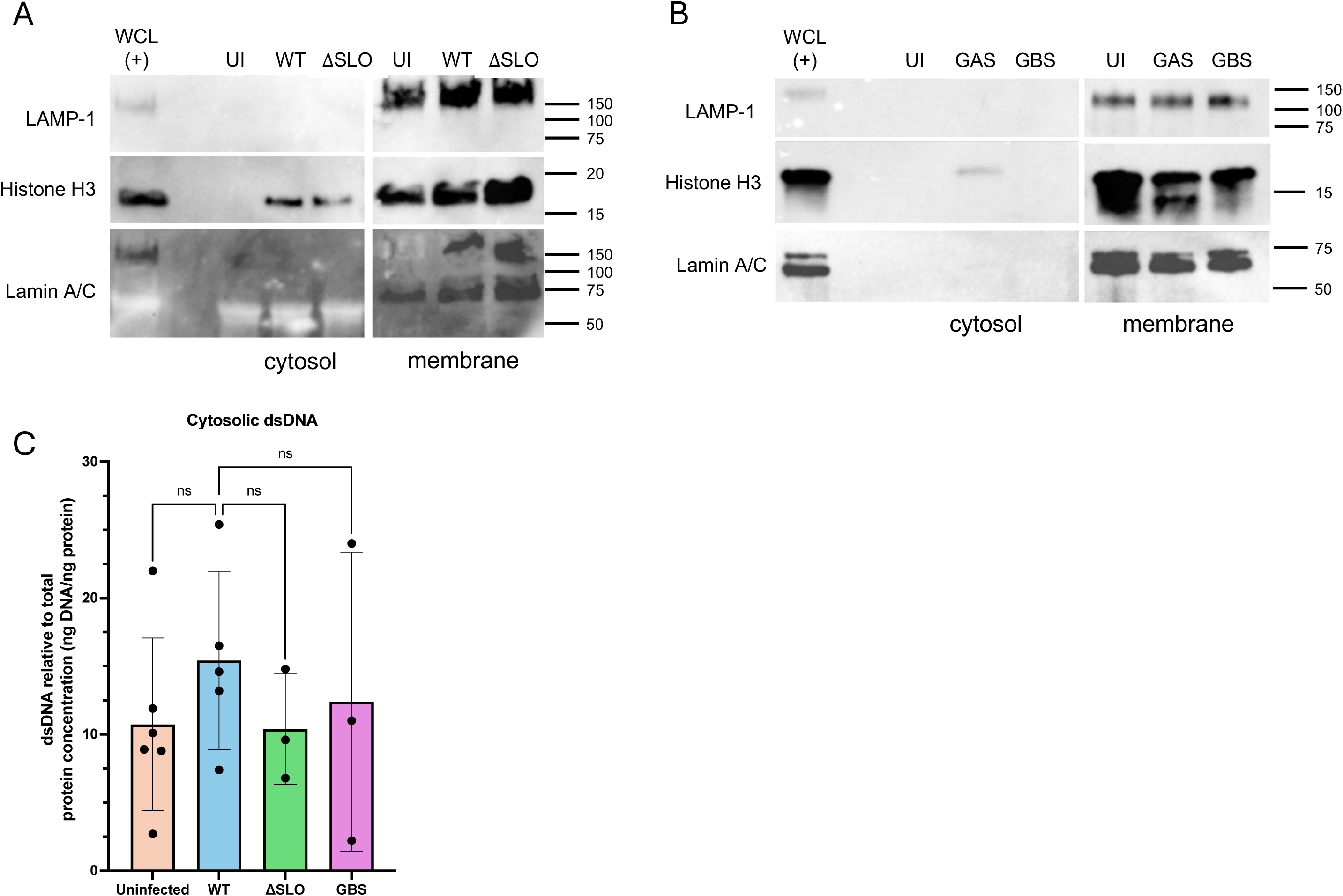
Histones are present in cytosolic preparations of GAS-infected cells. Cytosolic fractions from uninfected (UI), WT GAS-infected and ΔSLO-infected (SLO) cells **(A)** or GAS-infected (GAS) and GBS-infected (GBS) cells **(B)** were probed with the indicated antibodies. Whole cell lysates (WCL) and membrane fractions were included as positive controls and molecular weights are indicated. Experiments were performed at least 3 independent times and representative blots are shown. **C)** Double stranded DNA (dsDNA) content in the indicated cytosolic fractions was assessed. DNA amount was normalized to the protein concentration of the sample. Experiments were performed in duplicate at least 3 independent times and data are represented as mean ± SD and analyzed by one-way ANOVA with Tukey’s multiple comparison (ns = not significant).

The presence of all histone subunits in our infected cytosolic fractions (Supp. Fig. 2) suggests that nuclear DNA could be released from GAS-infected cells. We next quantified DNA in our cytosolic preparations. Although a low amount of DNA was detectable, there was no significant difference in the amount of DNA in cytosolic preparations from uninfected cells compared with GAS- or GBS-infected cells (Fig. 5C). These results are in agreement with previous findings that GAS does not trigger the cytosolic double-stranded DNA sensor cGAS in macrophages (38). Thus, GAS infection causes histones to be present in the cytosol in the absence of DNA, though the mechanism for their presence is unknown.

## Discussion

The human-specific pathogen GAS has co-evolved with the human immune system and has multiple strategies for survival within the host (39, 40). Although typically considered an extracellular bacterial pathogen, GAS can survive in macrophages, altering their functions and immune activity, and using them as shelters against antibiotic treatment (12,15,16). Modulating the inflammatory response of macrophages can further promote GAS survival by eliminating microbial competitors and recruiting T cells to the site of infection (39).

In this work, we determined which proteins have altered expression in the macrophage cytosol during GAS infection, both for regularly expressed cytosolic proteins and proteins that could be introduced through GAS permeabilization of phagosomes such as lysosomal proteins. Although we identified many proteins in our cytosolic fractions as expected, our datasets surprisingly revealed a relatively small number of proteins differentially expressed between infected and non-infected cells (Fig. 3). Furthermore, there is a larger difference in overall protein expression in ΔSLO- infected cells compared with WT-infected and uninfected cells (Fig. 3). These differences in part reflect the size and extent of phagosomal perforation mediated by the different strains of bacteria, as well as the consequences of releasing different proteins into the cytosol (15,16). The limited number of GAS proteins detected in our proteomics analysis gives confidence in our method that we were not identifying contaminants (Fig. 3, Supp. Tables 1-3). On the other hand, the proteins we did detect (e.g. M1, GAPDH, SLO, NADase) are secreted proteins that often are studied as vaccine candidates in part due to the propensity of GAS to produce them (41–43). Other GAS proteins such as pyruvate dehydrogenase may be playing a role in modifying the macrophage environment to dampen the inflammatory response (44), but we did not detect those proteins in our cytosolic fractions. Whether such bacterial proteins were below the limit of our detection or play an indirect role in controlling the host response is unknown and should be further studied.

In this study, we used IL-1β response to monitor release of phagosomal proteins into the cytosol since the NLRP3 inflammasome is a sensor for intracellular danger (22). We not only confirmed that M1 protein can stimulate the NLRP3 inflammasome response, but our data support that M1 protein is leaked from phagosomes into the cytosol in WT-infected cells (Fig. 4), providing a mechanism by which M1 can trigger an NLRP3 inflammasome response during natural infection. In WT infections, release of both M1 and SLO into the cytosol could contribute to the IL-1β response we measured (Fig. 2, 4, Supp. Table 2). Surprisingly, cytosolic fractions from ΔSLO-infected cells also elicited a strong IL-1β response (Fig. 2). In both WT- and ΔSLO-infected cytosol, we found cathepsin B (Supp. Table 2), which has been shown to process pro-IL-1β (19).

However, using the cathepsin B-specific inhibitor, we showed that this is likely not contributing to the IL-1β response we observe (Fig. 4). In previous work, we observed that ΔSLO bacteria also permeabilize phagosomes using CD63/LAMP-3 (16). This likely produces smaller pores than SLO, preventing larger proteins such as M1 from being released, but allowing other bacterial proteins to escape (16). We did find the GAS cysteine protease SpeB in the cytosol of ΔSLO-infected cells (Supp. Table 2), which has been shown to directly process pro-IL-1β (23), and may explain the ability of the ΔSLO mutant to elicit the IL-1β response.

A surprising finding from our data was the presence of histones in the cytosol (Fig. 3, 5). GAS is a well-known inducer of neutrophil extracellular traps (NETs), where extracellularly released histone-associated DNA mixes with antimicrobial compounds to trap and kill pathogens (45,46). Release of extracellular histone-associated DNA by macrophages in bacterial infection has been documented (23,47). Extracellular trap formation requires the breakdown or pore-formation in the nuclear envelope, and such damage could be mediated by SLO, the NLRP3 inflammasome, or gasdermin D, which can be activated by the GAS enzyme SpeB (23,26,47,48). GAS infection does not cause DNA to be present in the cytosol of cells, preventing detection by the double-stranded DNA sensor cGAS (Fig. 5)(38). However, DNA could be degraded by GAS DNase (45), leaving histones in the cytosol. Although we performed a preliminary analysis for nuclear modifications of histones such as methylation, we could not determine whether histones in our cytosolic fractions were of nuclear origin due to low concentration (Fig. 5), thus the origin and mechanism of histone release in GAS infection remain to be determined. Histone proteins have antimicrobial activity against GAS, and inflammatory capacity when released extracellularly, which may be responsible for sepsis during severe invasive infections (46,49,50) and thus warrants further study.

Inflammation during GAS infection can have varying consequences depending on the anatomic location. Inflammation in the nasopharynx, for example, is beneficial for both eliciting a strong protective immune response and promoting GAS survival, while inflammation in soft tissues such as skin can result in the severe pathologies associated with iGAS infection (21,23,39). Furthermore, persistent IL-1ꞵ activation as a result of GAS infection has been linked with ARF and rheumatic heart disease (51). However, caution must be exercised in using anti-inflammatory therapies, as IL-1 suppressants such as Anakinra are associated with increased iGAS outcomes (21). Our work demonstrates that multiple proteins, contributed both by the bacteria and the host, trigger the IL-1ꞵ response as a result of phagosomal permeabilization and the failure of macrophages to degrade and kill GAS. Therapies for GAS infection should therefore be aimed at the restoration of phagosome function, rather than the subsequent inflammatory response, to allow individuals to mount a proper protective immune response.

### Author Contributions

CO, CD, and JY, III contributed to conception and design of the study. All authors performed experiments and analyzed data. CO wrote the first draft of the manuscript. All authors contributed to manuscript revision, read, and approved the submitted version.

## Acknowledgments

We thank Isabelle Yuen, Henry Franscioni, Mitchell Johnson, Kyrlia Young and Tylor Lee for contributions to experiments, Shana Goffredi for the use of the Qubit fluorometer, and Clara Neville, Tylor Lee and Shana Goffredi for helpful discussions.

## Data availability

Mass spectrometry spectra and analysis parameters are available in the MassIVE database, accession number PXD069795.

https://massive.ucsd.edu/ProteoSAFe/dataset.jsp?task=bdb7b450c3d54c598461ad6e23ce44be

Reviewers only

Username: MSV000099561_reviewer

Password : 9dvH8k8yNj2iTKsM

## Packages

Stekhoven DJ (2022). *missForest: Nonparametric Missing Value Imputation using Random Forest*. R package version 1.5.

## Supplemental data

### Supplemental methods

Cells co-cultured with cell fractions were pre-treated with lipopolysaccharide from Escherichia coli K12 (LPS) for signal 1 enhancement and incubated for 24 hours at 37°C, 5% CO2.

**Supp. Figure 1:**
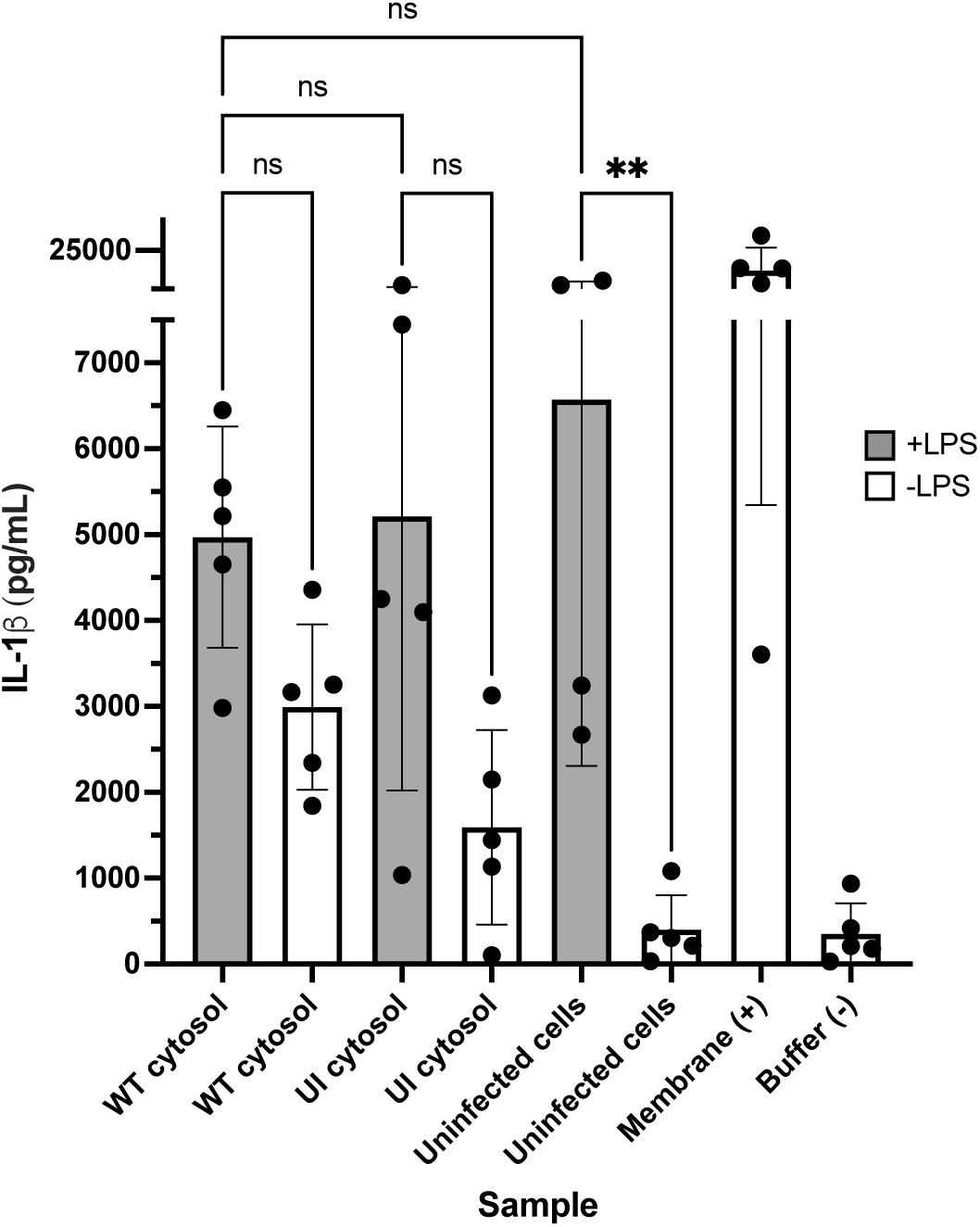
THP-1 cells pre-stimulated with LPS secrete a high basal level of IL-1b. THP-1 cells were incubated with 10ng/mL *E. coli* LPS (+LPS) or media only (-LPS) 24 hours before incubation with cytosolic fractions from WT GAS-infected (WT) or uninfected (UI) cells. THP-1 cells incubated in media only (uninfected cells), membrane fractions or fractionation buffer were included as controls. Data are represented as mean ± SD and analyzed by one-way ANOVA with Tukey’s multiple comparison (*p<0.05, **p<0.01, ns = not significant). A significant effect of LPS was found by two-way ANOVA (p<0.0001, data not shown).

**Supp. Figure 2:**
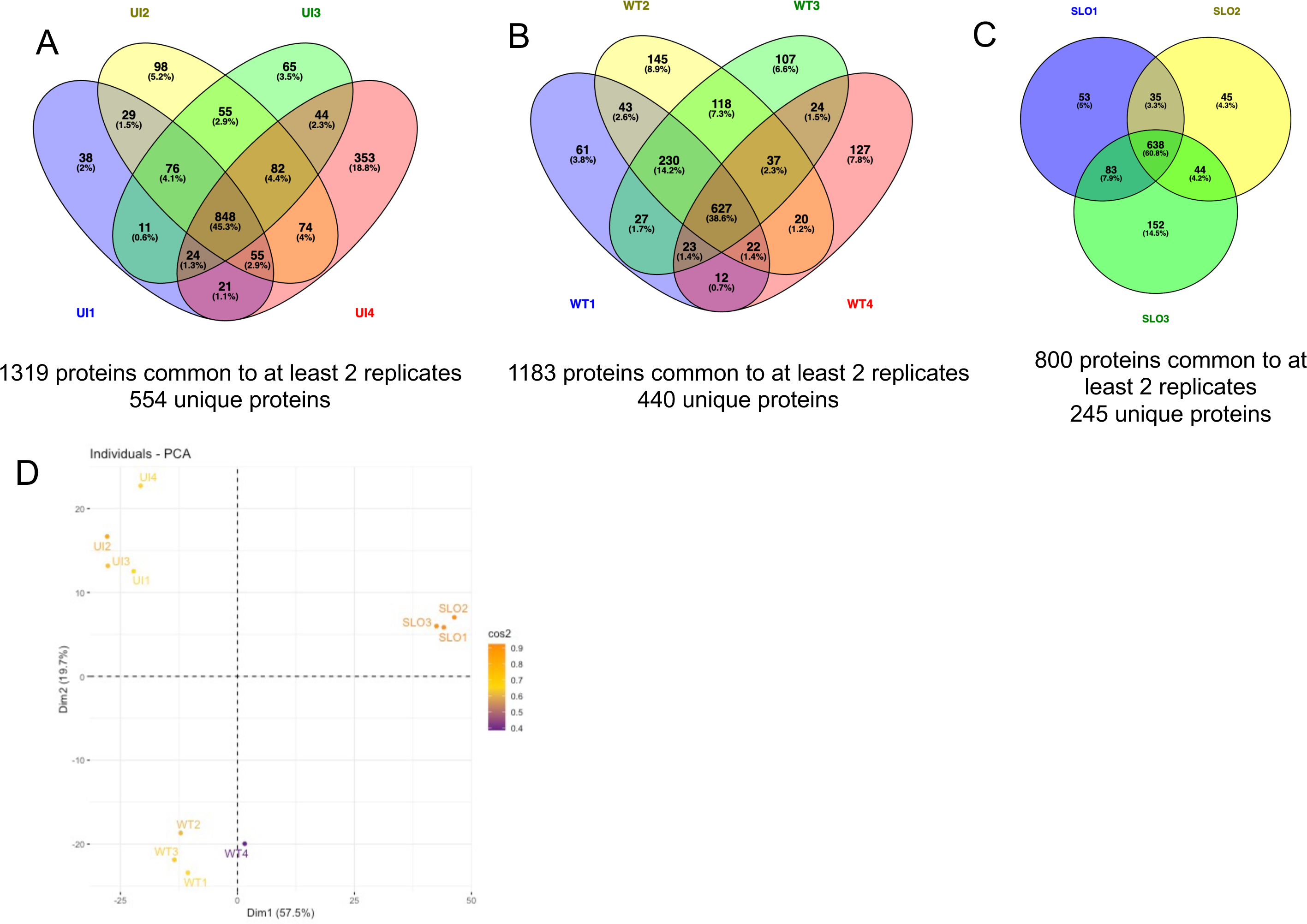
Number of proteins identified in each independent preparation of cytosolic fractions of **A)** uninfected (UI), **B)** WT-infected (WT) and **C)** τιSLO-infected (SLO) cells. Number of proteins common to at least 2 replicates and number of unique (low abundance) proteins are indicated. Images were generated with Venny v.2.1 (https://bioinfogp.cnb.csic.es/tools/venny/). **D)** PCA analysis of all individual preparations.

**Supp. Figure 3:**
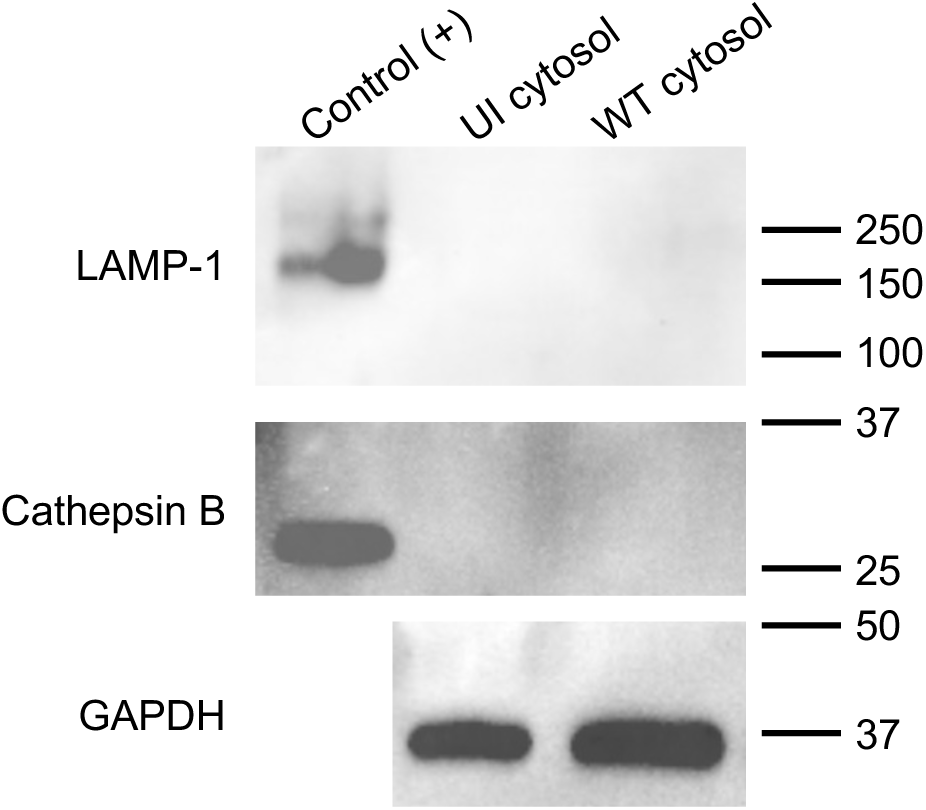
Cathepsin B is not detectable by Western blot in cytosolic fractions. Cytosolic proteins >30kD were isolated from uninfected (UI) or WT-infected THP-1 cells and probed for LAMP-1, Cathepsin B or GAPDH (cytosolic protein). The membrane fraction from WT-infected THP-1 cells was included as a control. Experiments were performed at least 3 independent times, and a representative blot is shown.\

**Supp. Table 1:** All data from the proteomics of cytosolic fractions collected from uninfected (UI), WT-infected (WT) and ΔSLO-infected (SLO) THP-1 macrophages.

**Supp. Table 2:** Selected proteomics data of unique proteins from cytosolic fractions collected from uninfected (UI), WT-infected (WT) and ΔSLO-infected (SLO) THP-1 macrophages.

**Supp. Table 3:** Pathway enrichment analyses of selected proteomics data of unique proteins from cytosolic fractions collected from uninfected (UI), WT-infected (WT) and ΔSLO-infected (SLO) THP-1 macrophages.

## References

1. Centers for Disease Control and Prevention. Active Bacterial Core Surveillance Report, Emerging Infections Program Network, Group A Streptococcus, 2023. [Internet]. 2023. Available from: www.cdc.gov/abcs/downloads/GAS_Surveillance_Report_2023.pdf

2. Gregory CJ, Okaro JO, Reingold A, Chai S, Herlihy R, Petit S, et al. Invasive Group A Streptococcal Infections in 10 US States. JAMA. 2025 May 6;333(17):1498–507.

3. Cobo-Vázquez E, Aguilera-Alonso D, Grandioso-Vas D, Gamell A, Rello-Saltor V, Oltra-Benavent M, et al. Sharp increase in the incidence and severity of invasive Streptococcus pyogenes infections in children after the COVID-19 pandemic (2019- 2023): A nationwide multicenter study. Int J Infect Dis. 2025 July 16;159:107982.

4. Colomina MS, Flamant A, Le Balle G, Cohen JF, Berthomieu L, Leteurtre S, et al. Severe Group A Streptococcus Infection among Children, France, 2022-2024. Emerg Infect Dis. 2025 Sept;31(9):1698–707.

5. Musumeci S, MacPhail A, Weisser Rohacek M, Erba A, Goldenberger D, Lang C, et al. Role of viral coinfection in post-pandemic invasive Group A streptococcal infections in adults, a nation-wide cohort study (iGASWISS). Eur J Clin Microbiol Infect Dis. 2025 Aug 5;

6. van Kempen EB, Tulling AJ, von Asmuth EGJ, van der Aa LB, Bijker EM, Bijlsma M, et al. Risk Factors for Severe Pediatric Invasive Group A Streptococcal Disease. JAMA Netw Open. 2025 Aug 1;8(8):e2527717.

7. Keller N, Andreoni F, Reiber C, Luethi-Schaller H, Schuepbach RA, Moch H, et al. Human Streptococcal Necrotizing Fasciitis Histopathology Mirrored in a Murine Model. The American Journal of Pathology. 2018 July 1;188(7):1517–23.

8. Avendaño-Ortiz J, Aguilera-Alonso D, Rodríguez CM, Oteo-Iglesias J, Lozano- Rodríguez R, López-Collazo E, et al. Plasma calprotectin as a severity biomarker in pediatric invasive *Streptococcus pyogenes* infections: insights from a multicenter immune profiling study during an outbreak. Journal of Microbiology, Immunology and Infection [Internet]. 2025 July 15 [cited 2025 Sept 8]; Available from: https://www.sciencedirect.com/science/article/pii/S1684118225001434

9. Bläckberg A, Svedevall S, Lundberg K, Nilson B, Kahn F, Rasmussen M. Time to Blood Culture Positivity: An Independent Predictor of Mortality in Streptococcus Pyogenes Bacteremia. Open Forum Infect Dis. 2022 June;9(6):ofac163.

10. Goldmann O, Lehne S, Medina E. Age-related susceptibility to Streptococcus pyogenes infection in mice: underlying immune dysfunction and strategy to enhance immunity. J Pathol. 2010 Apr;220(5):521–9.

11. Mishalian I, Ordan M, Peled A, Maly A, Eichenbaum MB, Ravins M, et al. Recruited macrophages control dissemination of group A Streptococcus from infected soft tissues. J Immunol. 2011 Dec 1;187(11):6022–31.

12. Thulin P, Johansson L, Low DE, Gan BS, Kotb M, McGeer A, et al. Viable group A streptococci in macrophages during acute soft tissue infection. PLoS Med. 2006 Mar;3(3):e53.

13. Anderson J, Imran S, Frost HR, Azzopardi KI, Jalali S, Novakovic B, et al. Immune signature of acute pharyngitis in a Streptococcus pyogenes human challenge trial. Nat Commun. 2022 Feb 9;13(1):769.

14. Nordenfelt P, Grinstein S, Björck L, Tapper H. V-ATPase-mediated phagosomal acidification is impaired by Streptococcus pyogenes through Mga-regulated surface proteins. Microbes Infect. 2012 Nov;14(14):1319–29.

15. Bastiat-Sempe B, Love JF, Lomayesva N, Wessels MR. Streptolysin O and NAD- glycohydrolase prevent phagolysosome acidification and promote group A Streptococcus survival in macrophages. mBio. 2014 Sept 16;5(5):e01690–01614.

16. Nishioka ST, Snipper J, Lee J, Schapiro J, Zhang RZ, Abe H, et al. Group A Streptococcus induces lysosomal dysfunction in THP-1 macrophages. Infect Immun. 2024 June 11;92(6):e0014124.

17. O’Neill AM, Thurston TLM, Holden DW. Cytosolic Replication of Group A Streptococcus in Human Macrophages. mBio. 2016 Apr 12;7(2):e00020–00016.

18. Hornung V, Bauernfeind F, Halle A, Samstad EO, Kono H, Rock KL, et al. Silica crystals and aluminum salts activate the NALP3 inflammasome through phagosomal destabilization. Nat Immunol. 2008 Aug;9(8):847–56.

19. Chevriaux A, Pilot T, Derangère V, Simonin H, Martine P, Chalmin F, et al. Cathepsin B Is Required for NLRP3 Inflammasome Activation in Macrophages, Through NLRP3 Interaction. Front Cell Dev Biol. 2020;8:167.

20. Harder J, Franchi L, Muñoz-Planillo R, Park JH, Reimer T, Núñez G. Activation of the Nlrp3 inflammasome by Streptococcus pyogenes requires streptolysin O and NF-kappa B activation but proceeds independently of TLR signaling and P2X7 receptor. J Immunol. 2009 Nov 1;183(9):5823–9.

21. LaRock CN, Todd J, LaRock DL, Olson J, O’Donoghue AJ, Robertson AAB, et al. IL- 1β is an innate immune sensor of microbial proteolysis. Sci Immunol. 2016 Aug;1(2):eaah3539.

22. Valderrama JA, Riestra AM, Gao NJ, LaRock CN, Gupta N, Ali SR, et al. Group A streptococcal M protein activates the NLRP3 inflammasome. Nat Microbiol. 2017 Oct;2(10):1425–34.

23. LaRock DL, Johnson AF, Wilde S, Sands JS, Monteiro MP, LaRock CN. Group A Streptococcus induces GSDMA-dependent pyroptosis in keratinocytes. Nature. 2022 May;605(7910):527–31.

24. Walker MJ, Hollands A, Sanderson-Smith ML, Cole JN, Kirk JK, Henningham A, et al. DNase Sda1 provides selection pressure for a switch to invasive group A streptococcal infection. Nat Med. 2007 Aug;13(8):981–5.

25. Chatellier S, Ihendyane N, Kansal RG, Khambaty F, Basma H, Norrby-Teglund A, et al. Genetic relatedness and superantigen expression in group A streptococcus serotype M1 isolates from patients with severe and nonsevere invasive diseases. Infect Immun. 2000 June;68(6):3523–34.

26. Timmer AM, Timmer JC, Pence MA, Hsu LC, Ghochani M, Frey TG, et al. Streptolysin O promotes group A Streptococcus immune evasion by accelerated macrophage apoptosis. J Biol Chem. 2009 Jan 9;284(2):862–71.

27. Lauth X, von Köckritz-Blickwede M, McNamara CW, Myskowski S, Zinkernagel AS, Beall B, et al. M1 protein allows Group A streptococcal survival in phagocyte extracellular traps through cathelicidin inhibition. J Innate Immun. 2009;1(3):202–14.

28. Wilson CB, Weaver WM. Comparative susceptibility of group B streptococci and Staphylococcus aureus to killing by oxygen metabolites. J Infect Dis. 1985 Aug;152(2):323–9.

29. Repnik U, Borg Distefano M, Speth MT, Ng MYW, Progida C, Hoflack B, et al. L- leucyl-L-leucine methyl ester does not release cysteine cathepsins to the cytosol but inactivates them in transiently permeabilized lysosomes. J Cell Sci. 2017 Sept 15;130(18):3124–40.

30. Aguilan JT, Kulej K, Sidoli S. Guide for protein fold change and p-value calculation for non-experts in proteomics. Mol Omics. 2020 Dec 1;16(6):573–82.

31. Stekhoven DJ, Bühlmann P. MissForest--non-parametric missing value imputation for mixed-type data. Bioinformatics. 2012 Jan 1;28(1):112–8.

32. Galili T, O’Callaghan A, Sidi J, Sievert C. heatmaply: an R package for creating interactive cluster heatmaps for online publishing. Bioinformatics. 2018 May 1;34(9):1600–2.

33. Ge SX, Jung D, Yao R. ShinyGO: a graphical gene-set enrichment tool for animals and plants. Bioinformatics. 2020 Apr 15;36(8):2628–9.

34. Richter J, Monteleone MM, Cork AJ, Barnett TC, Nizet V, Brouwer S, et al. Streptolysins are the primary inflammasome activators in macrophages during Streptococcus pyogenes infection. Immunology & Cell Biology. 2021;99(10):1040– 52.

35. Tsai CM, Riestra AM, Ali SR, Fong JJ, Liu JZ, Hughes G, et al. Siglec-14 Enhances NLRP3-Inflammasome Activation in Macrophages. J Innate Immun. 2019 Dec 5;12(4):333–43.

36. Kang MJ, Jo SG, Kim DJ, Park JH. NLRP3 inflammasome mediates interleukin-1β production in immune cells in response to Acinetobacter baumannii and contributes to pulmonary inflammation in mice. Immunology. 2017;150(4):495–505.

37. Buttle DJ, Murata M, Knight CG, Barrett AJ. CA074 methyl ester: a proinhibitor for intracellular cathepsin B. Arch Biochem Biophys. 1992 Dec;299(2):377–80.

38. Movert E, Lienard J, Valfridsson C, Nordström T, Johansson-Lindbom B, Carlsson F. Streptococcal M protein promotes IL-10 production by cGAS-independent activation of the STING signaling pathway. PLoS Pathog. 2018 Mar;14(3):e1006969.

39. Guerra S, LaRock C. Group A Streptococcus interactions with the host across time and space. Curr Opin Microbiol. 2024 Feb;77:102420.

40. Bergsten H, Nizet V. The intricate pathogenicity of Group A Streptococcus: A comprehensive update. Virulence. 2024 Dec 31;15(1):2412745.

41. Osowicki J, Frost HR, Azzopardi KI, Whitcombe AL, McGregor R, Carlton LH, et al. Streptococcus pyogenes pharyngitis elicits diverse antibody responses to key vaccine antigens influenced by the imprint of past infections. Nat Commun. 2024 Dec 3;15(1):10506.

42. Bensi G, Mora M, Tuscano G, Biagini M, Chiarot E, Bombaci M, et al. Multi high- throughput approach for highly selective identification of vaccine candidates: the Group A Streptococcus case. Mol Cell Proteomics. 2012 June;11(6):M111.015693.

43. Jin H, Agarwal S, Agarwal S, Pancholi V. Surface export of GAPDH/SDH, a glycolytic enzyme, is essential for Streptococcus pyogenes virulence. mBio. 2011;2(3):e00068–00011.

44. Xu W, Bradstreet TR, Zou Z, Hickerson S, Zhou Y, He H, et al. Reprogramming aerobic metabolism mitigates Streptococcus pyogenes tissue damage in a mouse necrotizing skin infection model. Nat Commun. 2025 Mar 15;16(1):2559.

45. Buchanan JT, Simpson AJ, Aziz RK, Liu GY, Kristian SA, Kotb M, et al. DNase expression allows the pathogen group A Streptococcus to escape killing in neutrophil extracellular traps. Curr Biol. 2006 Feb 21;16(4):396–400.

46. Döhrmann S, LaRock CN, Anderson EL, Cole JN, Ryali B, Stewart C, et al. Group A Streptococcal M1 Protein Provides Resistance against the Antimicrobial Activity of Histones. Sci Rep. 2017 Feb 21;7:43039.

47. Lee Y, Brenner M, Aziz M, Wang P. Molecular and subcellular mechanisms of vital macrophage extracellular trap formation. Front Immunol [Internet]. 2025 July 31 [cited 2025 Sept 15];16. Available from: https://www.frontiersin.org/journals/immunology/articles/10.3389/fimmu.2025.16084 28/full

48. Münzer P, Negro R, Fukui S, di Meglio L, Aymonnier K, Chu L, et al. NLRP3 Inflammasome Assembly in Neutrophils Is Supported by PAD4 and Promotes NETosis Under Sterile Conditions. Front Immunol. 2021;12:683803.

49. Xu J, Zhang X, Pelayo R, Monestier M, Ammollo CT, Semeraro F, et al. Extracellular histones are major mediators of death in sepsis. Nat Med. 2009 Nov;15(11):1318– 21.

50. Westman J, Chakrakodi B, Snäll J, Mörgelin M, Bruun Madsen M, Hyldegaard O, et al. Protein SIC Secreted from Streptococcus pyogenes Forms Complexes with Extracellular Histones That Boost Cytokine Production. Front Immunol. 2018;9:236.

51. Kim ML, Martin WJ, Minigo G, Keeble JL, Garnham AL, Pacini G, et al. Dysregulated IL-1β-GM-CSF Axis in Acute Rheumatic Fever That Is Limited by Hydroxychloroquine. Circulation. 2018 Dec 4;138(23):2648–61.

## References

52. Oliveros JC. Venny. An interactive tool for comparing lists with Venn’s diagrams. https://bioinfogp.cnb.csic.es/tools/venny/index.html [Internet]. 2007.

